# NetGO 3.0: Protein Language Model Improves Large-scale Functional Annotations

**DOI:** 10.1101/2022.12.05.519073

**Authors:** Shaojun Wang, Ronghui You, Yunjia Liu, Yi Xiong, Shanfeng Zhu

## Abstract

As one of the state-of-the-art automated function prediction (AFP) methods, NetGO 2.0 integrates multi-source information to improve the performance. However, it mainly utilizes the proteins with experimentally supported functional annotations without leveraging valuable information from a vast number of unannotated proteins. Recently, protein language models have been proposed to learn informative representations (e.g., Evolutionary Scale Modelling (ESM)-1b embedding) from protein sequences based on self-supervision. We represent each protein by ESM-1b and use logistic regression (LR) to train a new model, LR-ESM, for AFP. The experimental results show that LR-ESM achieves comparable performance with the best-performing component of NetGO 2.0. Therefore, by incorporating LR-ESM into NetGO 2.0, we develop NetGO 3.0 to improve the performance of AFP extensively. NetGO 3.0 is freely accessible at https://dmiip.sjtu.edu.cn/ng3.0.

## Introduction

Proteins are complex molecules that play essential roles in various biological activities. To understand the underlying mechanism of an organism as a physical system, annotating the functions of proteins is a crucial task. Gene Ontology came into being in 1998 to describe varying levels of functional information on gene/RNA/protein, which contains three domains: Molecular Function Ontology (MFO), Biological Process Ontology (BPO), and Cellular Component Ontology (CCO) with over 40 000 terms [1]. As of Nov 2022, the number of raw protein sequences is more than 230 million in UniProtKB, but less than 0.1% of them have experimental annotations [2]. It is thus desirable to develop high-performing computational methods to achieve automated function prediction without costly experiments [3].

AFP is a large-scale multi-label classification problem where multiple related GO terms are assigned to a target protein. In the last few years, several high-performing web servers have been developed for AFP, such as INGA 2.0 [4], DeepGOWeb [5], MetaGO [6], and QAUST [7]. Under the learning to rank (LTR) framework [8], GOLabeler [9], NetGO [10], and NetGO 2.0 [11] achieved a state-of-the-art performance in the recent community-wide Critical Assessment of Functional Annotation (CAFA) [3]. Specifically, NetGO 2.0 integrates protein information from different sources to encode proteins in a computer-understandable way, such as sequences, protein domains, protein-protein interaction networks, and scientific literature. However, it does not leverage valuable information from unannotated proteins (>99.9% of all known proteins).

Recently, the idea of pre-training in natural language processing [12] has been extended to build protein language models using self-supervised learning on millions of sequences [13–15]. Most protein language models predict the masked or next amino acid within a sequence and generate protein embeddings that can generalize across downstream tasks (more details shown in Supplementary Materials). Some recent studies have explored protein language models for AFP [16, 17]. However, they have a common limitation: less frequent GO terms (e.g., having less than 40 annotated proteins) are excluded in the evaluation, which accounts for around 75% of total annotations in the CAFA setting. In this work, we predict the associations between proteins and each GO term based on ESM-1b embeddings, which were trained on over 250 million protein sequences [13]. Our experimental results show that the learned representations are helpful to AFP. Therefore, we develop NetGO 3.0 by incorporating ESM-1b embeddings in order to improve the performance extensively, which highlights the predictive power of the protein language model for AFP.

## Materials and methods

### Protein language models

A challenging problem is figuring out how to represent protein sequences as fixed-length vectors that capture the realistic sequence-function relationship. Traditional methods rely on a holistic understanding of protein properties. Recently, protein language models have provided a solution that interprets protein sequences as sentences and amino acids as words to extract fundamental features of a protein with rich and systematic information. Protein language models train nonlinear neural networks with an unsupervised objective on a large-scale dataset of protein sequences [13–15, 21].

Generally, protein language models apply deep learning models such as recurrent neural networks (RNN) and Transformer to achieve statistical embeddings of proteins from tremendous sequences. UniRep represented protein sequences as fixed-length vectors by LSTM with ~24 million sequences [15]. TAPE distilled protein properties from sequences by semi-supervised learning based on ResNet, LSTM, and Transformer; then evaluated their performance on five biologically relevant tasks [21]. Moreover, a multi-task learning framework was recently proposed to incorporate structural information (e.g., contact maps and structural similarity prediction tasks) to enrich protein language models [22]. Furthermore, researchers applied protein language models to study protein molecular function prediction[17]. UDSMProt put forward a task-agnostic representation for proteins and achieved good performance on protein-level prediction tasks, namely enzyme class prediction and GO prediction [16]. However, both methods should have considered less frequent GO terms.

In this paper, a new component LR-ESM in NetGO 3.0 is proposed to utilize ESM-1b, a 34-layer Transformer-based model trained on UniPAC database with 250M protein sequences and 650M parameters, to generate protein-level representations by average pooling across all residue-level embeddings [13].

### Implementation

NetGO 2.0 integrates seven component methods, which are Naive, BLAST-KNN, LR-3mer, LR-InterPro, NetKNN, LR-Text, and Seq-RNN. We replace Seq-RNN with LR-ESM in NetGO 3.0, which makes function prediction based on a protein language model. Specifically, LR-ESM utilizes ESM-1b, a 34-layer Transformer-based model trained on the UniPac database with 250 million sequences [13], to generate protein embeddings and complete prediction. As ESM-1b has a limitation of protein sequence length, we keep the first 1000 amino acids for those protein sequences longer than 1024. We then use ESM-1b to encode each amino acid as an embedding of size 1280 for a target protein. To obtain the protein-level embedding, we apply the operation of average pooling on all amino acid positions, which comprehensively collects information from sequence data alone. Finally, LR-ESM utilizes protein embeddings as input to train logistic regression classifiers and estimates the association between target proteins and each GO term.

### Benchmark datasets

As NetGO 2.0 collected the data following the setting of CAFA, we utilized the same benchmark dataset to evaluate the performance of NetGO 3.0 and the competing methods. Table S1 in supplementary materials reports the number of proteins in the benchmark dataset.

To take advantage of the latest annotation data, we collect sequences and GO terms before January 2022 from UniProt [2], GOA [23] and GO [1]. Similarly, we train and update our model on the new dataset by following the standard protocols of NetGO 2.0 [11].

1. Training data: all experimental annotation data before January 2020.
2. Validation data: all experimental *no-knowledge* and *limited-knowledge* proteins annotated from January 2020 to December 2020.
3. Testing data: all experimental *no-knowledge* proteins between January 2021 and December 2021.

More details for the new dataset are listed in Section S2 and Table S2 of supplementary materials.

## Results

We compare the performance of NetGO 3.0 with the competing methods on the benchmark dataset from NetGO 2.0. The performance is evaluated by area under the precision-recall curve (AUPR) and two standard metrics in CAFA, *F_max_* and *S_min_*. The definitions of three metrics are given in Section S1 of supplementary materials.

### Performance comparisons of NetGO 3.0 with its component methods and competing methods

**Table 1** illustrates the test results for NetGO 3.0, NetGO 2.0, GOLabeler, DeepGOWeb, and the component methods of NetGO 3.0. GOLabeler and NetGO 2.0 achieved top performance in CAFA3 and CAFA4, respectively [9, 11]. DeepGOWeb provided an accurate prediction for protein function by deep learning [5]. The underlined numbers in the upper table imply the best performance for component methods.

**Table 1.**
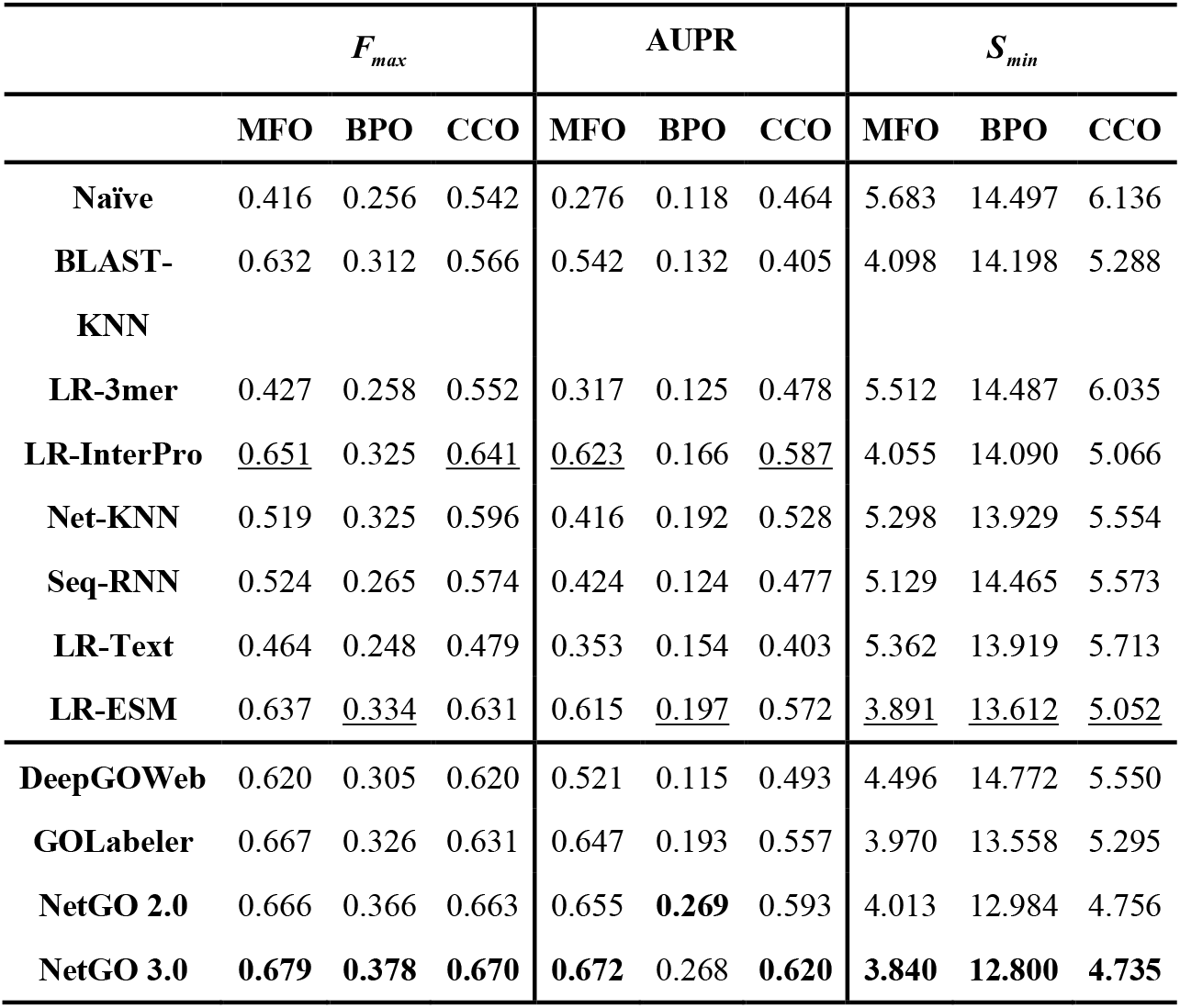
Performance of NetGO 3.0 with its components and competing methods on the test set.

We select Naive, BLAST-KNN, and Seq-RNN [11] from NetGO 2.0 as three baseline methods. The Naive method annotates each pair of protein and GO term with a score that equals the probability of the term appearing in the training data. BLAST-KNN assigns a protein with GO terms based on annotations of its top BLAST hits [9]. While the first two are component methods inherited from both NetGO and NetGO2.0, Seq-RNN is a new component of NetGO2.0, which was designed to extract the deep representation of a protein sequence [11]. As shown in **Figure 1**, LR-ESM outperforms baseline methods on all three GO domains. As a replacement for Seq-RNN, LR-ESM achieves a better performance. Specifically, in terms of *F_max_*, LR-ESM achieves 21.6%, 31.3%, and 7.5% improvements over Seq-RNN on MFO, BPO, and CCO, respectively, which indicates the effectiveness of ESM-1b for AFP. Moreover, LR-ESM and LR-InterPro show comparable performance in all three GO domains. Note that, in terms of *S_min_*, LR-ESM outperforms all other component methods and even achieves a better performance on MFO than NetGO 2.0. Therefore, it is reasonable to construct a more robust model by incorporating LR-ESM into NetGO 2.0.

**Figure 1.**
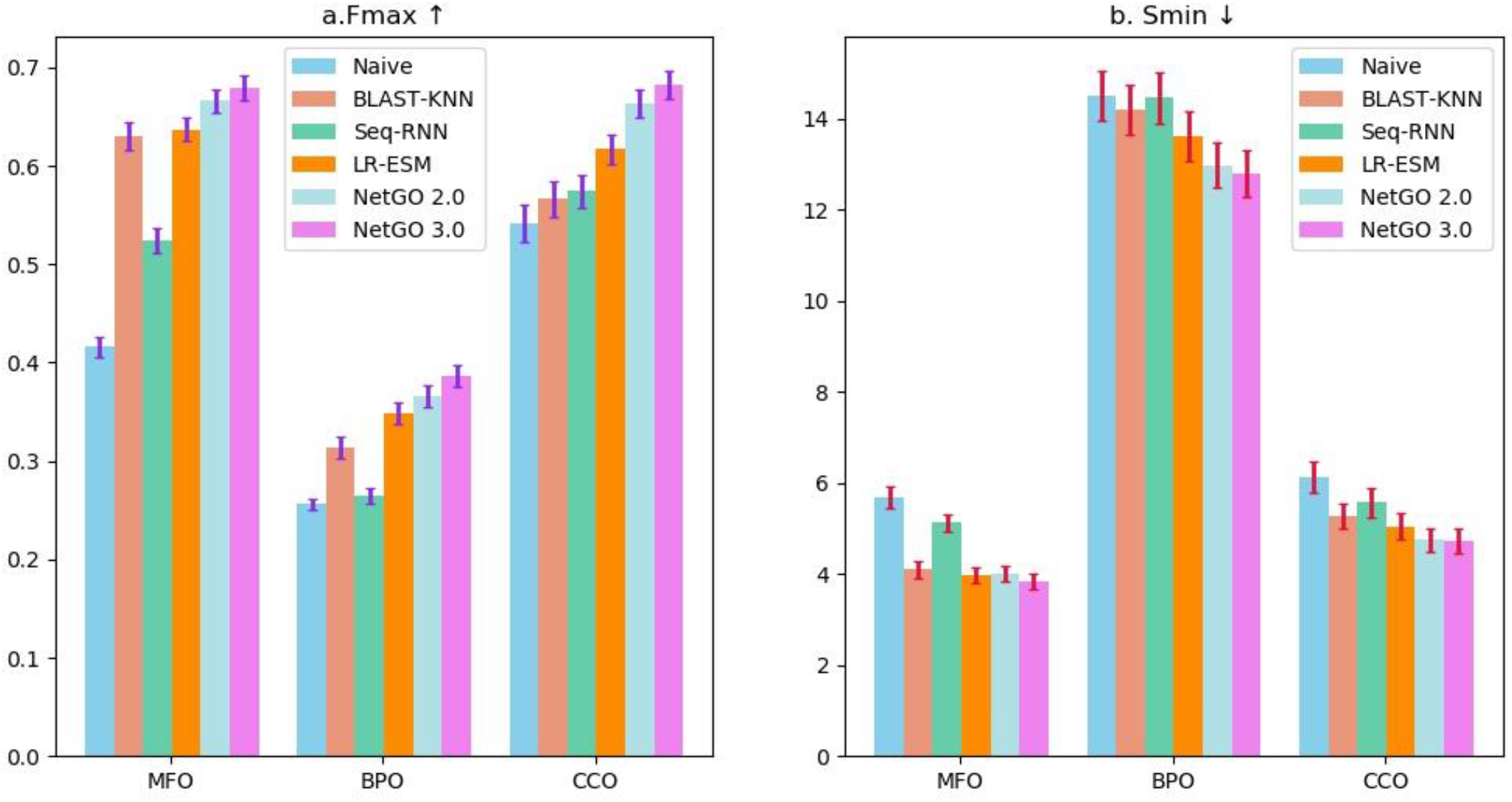
The performance comparison on *F_max_* and *S_min_*. The performance of Naive, BLAST-KNN, SeqRNN, LR-ESM, NetGO 2.0, and NetGO 3.0 on the benchmark dataset of NetGO 2.0 over three GO domains. The arrows imply higher values for *F_max_* and lower for *S_min_* over three GO domains. The bars denote the confidence intervals (95%) calculated by bootstrapping with 100 iterations as in CAFA.

Furthermore, we compare NetGO 3.0 with several high-performing methods in CAFA. In the lower part of Table 1, we highlight the best performance in bold among these methods. The experimental results show that NetGO 3.0 has achieved a more superior performance than the competing methods. In terms of *F_max_* and *S_max_*, NetGO 3.0 achieves a better performance in all three GO domains. For example, NetGO 3.0 achieved the highest *F_max_* of 0.378 in BPO, which was 3.3% and 16.0% improvements over NetGO 2.0 (0.366) and GOLabeler (0.326), respectively. Finally, we calculated the confidence interval (95%) for each method by bootstrapping with 100 iterations on the test set, as shown in Figure 1. The results demonstrate that NetGO 3.0 can benefit from protein language models with deep dense embeddings.

To better illustrate the strength of NetGO 3.0, we draw a Venn diagram in **Figure 2** to show the overlaps and differences among the prediction results of GOLabeler, DeepGOWeb, and NetGO 3.0. There are three main findings: (i) Although each method can predict distinct GO terms, the prediction results of the three methods are overlapped substantially, especially in CCO. Specifically, there are 6.96 GO terms assigned to one protein on average that are predicted by all three methods in CCO, which accounts for 62.5%, 70.1%, and 77.3% in the prediction results of DeepGOWeb (11.14), GOLabeler (9.84), and NetGO 3.0 (9.00), respectively. (ii) DeepGOWeb predicted more GO terms but achieved lower performance than the other two methods, indicating that false positive GO terms are common in the prediction results. For example, DeeGOWeb predicted 21.34 distinct GO terms and achieved the lowest *F_max_* of 0.305 in BPO, which suggests that most of its predicted GO terms are incorrect. (iii) Compared with MFO and CCO, NetGO 3.0 and GOLabeler differs significantly in predicting GO terms in BPO. Specifically, in terms of BPO, although the 15.42 GO terms predicted by the two methods are consistent, the number of distinct GO terms predicted by NetGO 3.0 and GOLabeler are 9.59 and 8.18, respectively. We note that NetGO 3.0 performed better than GOLabeler in BPO in terms of *F_max_*, where NetGO 3.0 (0.378) achieved a 16.0% improvement compared to GOLabeler (0.326). It demonstrates that NetGO 3.0 is more accurate and can predict more true positive terms for query proteins.

**Figure 2.**
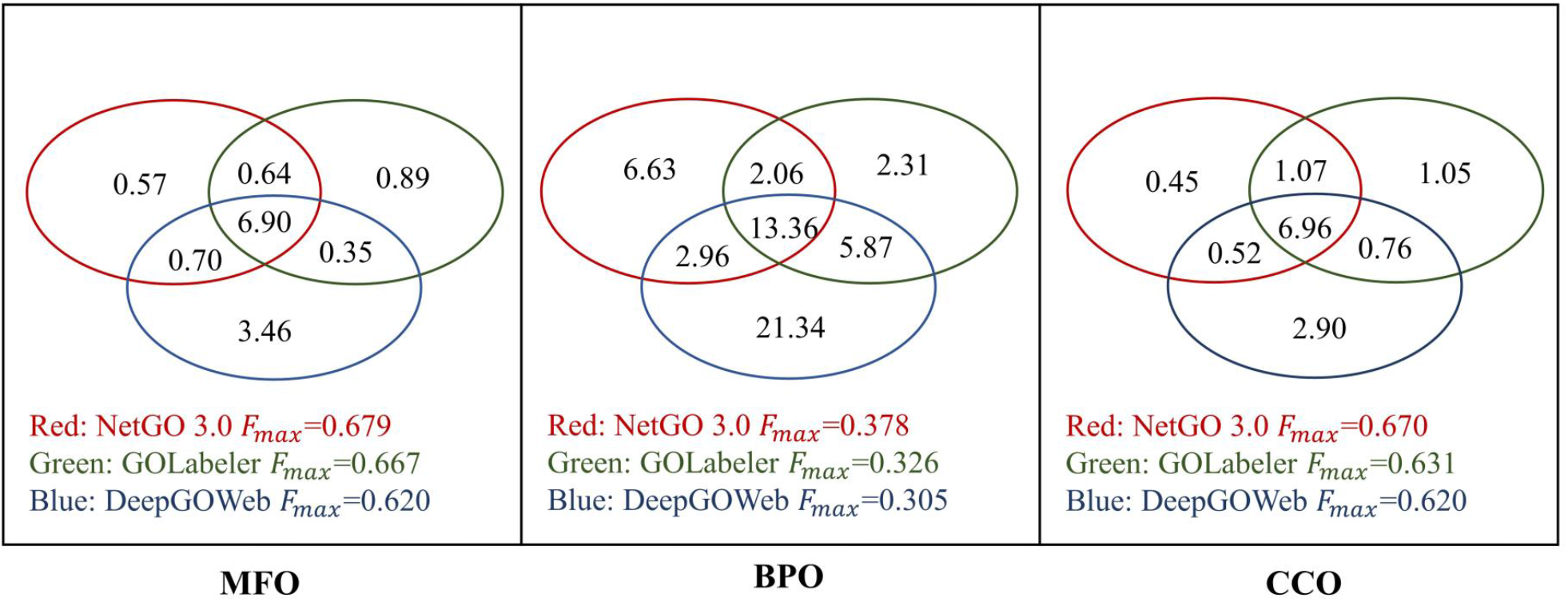
The overlap and difference among the predicted GO terms of GOLabeler, DeepGOWeb, and NetGO 3.0. The Venn diagram depicts the overlap and difference among the predicted GO terms of GOLabeler, DeepGOWeb, and NetGO 3.0. The figures in the graph represent the average number of predicted GO terms over test proteins in three methods.

### Performance on specific species (HUMAN and MOUSE)

Species-specific analyses are helpful for researchers to study a certain species. Here, we explore the performance of different AFP methods over two model species, HUMAN and MOUSE. **Tables 2** and **3** show the performance of NetGO 3.0 and the competing methods for proteins in HUMAN and MOUSE. According to the performance of the two species, we can see that all methods obtain a better prediction performance on HUMAN in comparison with MOUSE. For example, LR-InterPro, LR-ESM, and NetGO 3.0 achieved higher AUPRs of 0.704, 0.690, and 0.730 over HUMAN proteins in MFO, whereas the three methods only achieved AUPRs of 0.601, 0.615, and 0.620 over MOUSE proteins. The annotation information for different species is from different databases, which may lead to the difference. Moreover, LR-ESM again achieves a similar performance as LR-InterPro in both species, which strongly demonstrates that features extracted by ESM-1b are as robust as InterProScan among many species.

**Table 2.**
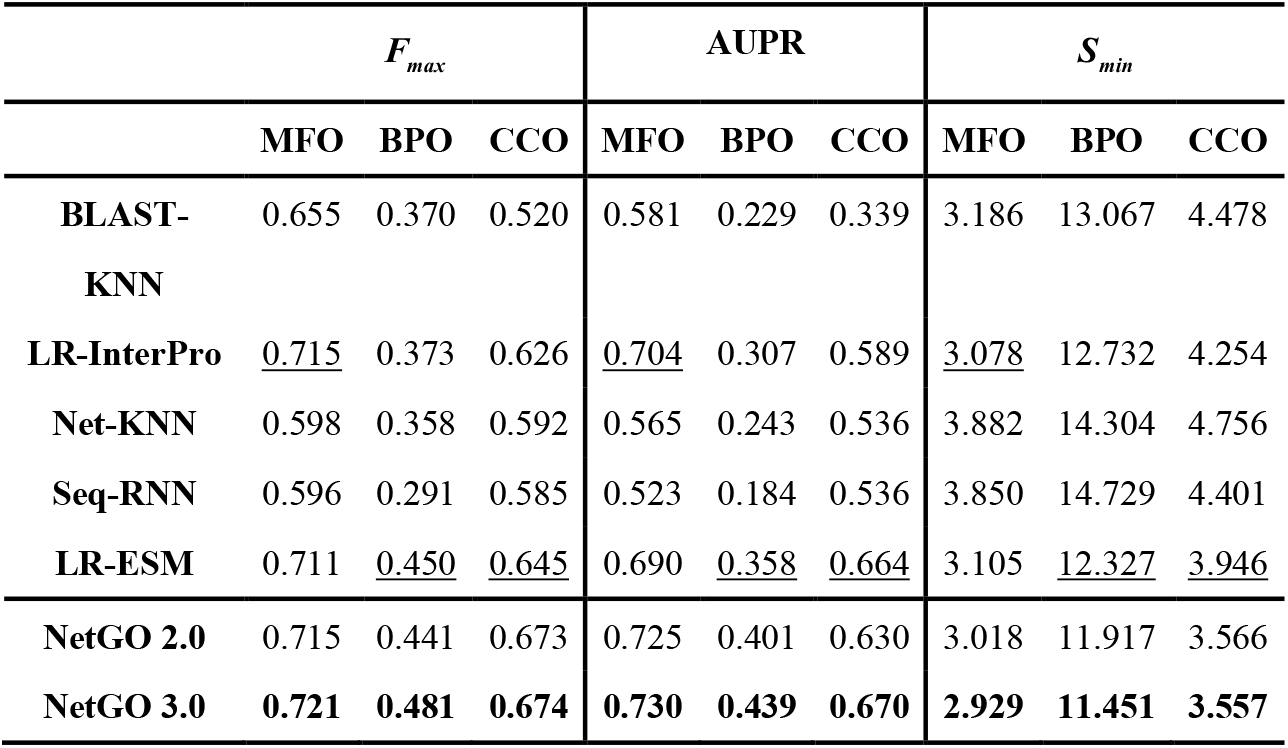
Performance comparisons for proteins in HUMAN.

**Table 3.** Performance comparisons for proteins in MOUSE.

| | $F_{max}$ | | | AUPR | | | $S_{min}$ | | |
| --- | --- | --- | --- | --- | --- | --- | --- | --- | --- |
|  | MFO | BPO | CCO | MFO | BPO | CCO | MFO | BPO | CCO |
| **BLAST-KNN** | 0.616 | 0.353 | 0.572 | 0.575 | 0.147 | 0.463 | 5.681 | 21.804 | 5.931 |
| **LR-InterPro** | 0.605 | 0.344 | <u>0.591</u> | 0.609 | <u>0.211</u> | <u>0.542</u> | 5.732 | 21.044 | <u>5.489</u> |
| **Net-KNN** | 0.408 | 0.341 | 0.566 | 0.253 | 0.199 | 0.502 | 8.080 | 21.309 | 6.002 |
| **Seq-RNN** | 0.520 | 0.265 | 0.537 | 0.373 | 0.106 | 0.462 | 6.787 | 22.795 | 5.993 |
| **LR-ESM** | <u>0.639</u> | <u>0.352</u> | 0.561 | <u>0.615</u> | 0.197 | 0.539 | <u>5.710</u> | <u>20.639</u> | 5.664 |
| **NetGO 2.0** | **0.649** | 0.420 | 0.617 | 0.618 | 0.315 | 0.557 | 5.683 | 19.572 | 5.563 |
| **NetGO 3.0** | **0.649** | **0.427** | **0.620** | **0.620** | **0.316** | **0.568** | **5.583** | **19.545** | **5.034** |

For HUMAN and MOUSE, NetGO 3.0 outperforms all competing methods in all three GO domains. Specifically, NetGO 3.0 performs better than NetGO 2.0 in HUMAN BPO prediction, which achieves 9.3% and 9.5% improvements in terms of *F_max_* and AUPR. Further, the results highlight the importance of source data and the effectiveness of the protein language model.

### Performance comparisons over groups categorized by the number of annotations per GO term

We divided GO terms in the test dataset into three groups according to the number of annotations per GO term: 10-30, 31-100, and >100. **Table 4** reports the M-AUPR computed in each group, where M-AUPR is GO term-centric by averaging AUPR on each GO term. LR-ESM outperforms other component methods in most cases, which indicates that ESM-1b embeddings are informative. Note that LR-ESM consistently ranks higher than LR-InterPro for three domains in the first group, especially for BPO, which obtains a 47.8% improvement. It proves that protein embeddings are effective with such a vast amount of training data for AFP.

**Table 4.** Performance comparisons over groups categorized by the number of annotations per GO term.

| Methods | MFO |  |  | BPO |  |  | CCO |  |  |
| --- | --- | --- | --- | --- | --- | --- | --- | --- | --- |
|  | 10-30 | 31-100 | >100 | 10-30 | 31-100 | >100 | 10-30 | 31-100 | >100 |
| **BLAST-KNN** | 0.628 | 0.497 | 0.614 | 0.197 | 0.131 | 0.224 | 0.265 | 0.291 | 0.528 |
| **LR-InterPro** | 0.618 | <u>0.562</u> | 0.634 | 0.209 | 0.138 | 0.224 | 0.231 | 0.307 | <u>0.589</u> |
| **Net-KNN** | 0.330 | 0.253 | 0.545 | 0.139 | 0.132 | 0.222 | 0.210 | 0.269 | 0.501 |
| **Seq-RNN** | 0.434 | 0.326 | 0.525 | 0.054 | 0.062 | 0.139 | 0.152 | 0.195 | 0.437 |
| **LR-ESM** | <u>0.642</u> | 0.516 | <u>0.658</u> | <u>0.307</u> | <u>0.154</u> | <u>0.242</u> | <u>0.333</u> | <u>0.342</u> | 0.572 |
| **NetGO 2.0** | 0.658 | 0.569 | 0.659 | 0.248 | 0.212 | 0.329 | 0.300 | 0.389 | 0.588 |
| **NetGO 3.0** | **0.675** | **0.571** | **0.665** | **0.250** | **0.213** | **0.335** | **0.386** | **0.422** | **0.636** |

NetGO 3.0 achieves the best results among all the methods in every group and domain, and the improvement over NetGO 2.0 is especially significant in CCO. Specifically, the advances made by NetGO 3.0 are 28.7%, 8.4%, and 8.2% for the three groups, respectively. Moreover, we collect the CCO terms in the second and third layers annotated with more than ten proteins in the test set. As shown in **Figure 3**, NetGO 3.0 achieves a better performance on most GO terms, which strongly suggests that ESM-1b is powerful for predicting protein functions about cellular components.

**Figure 3.**
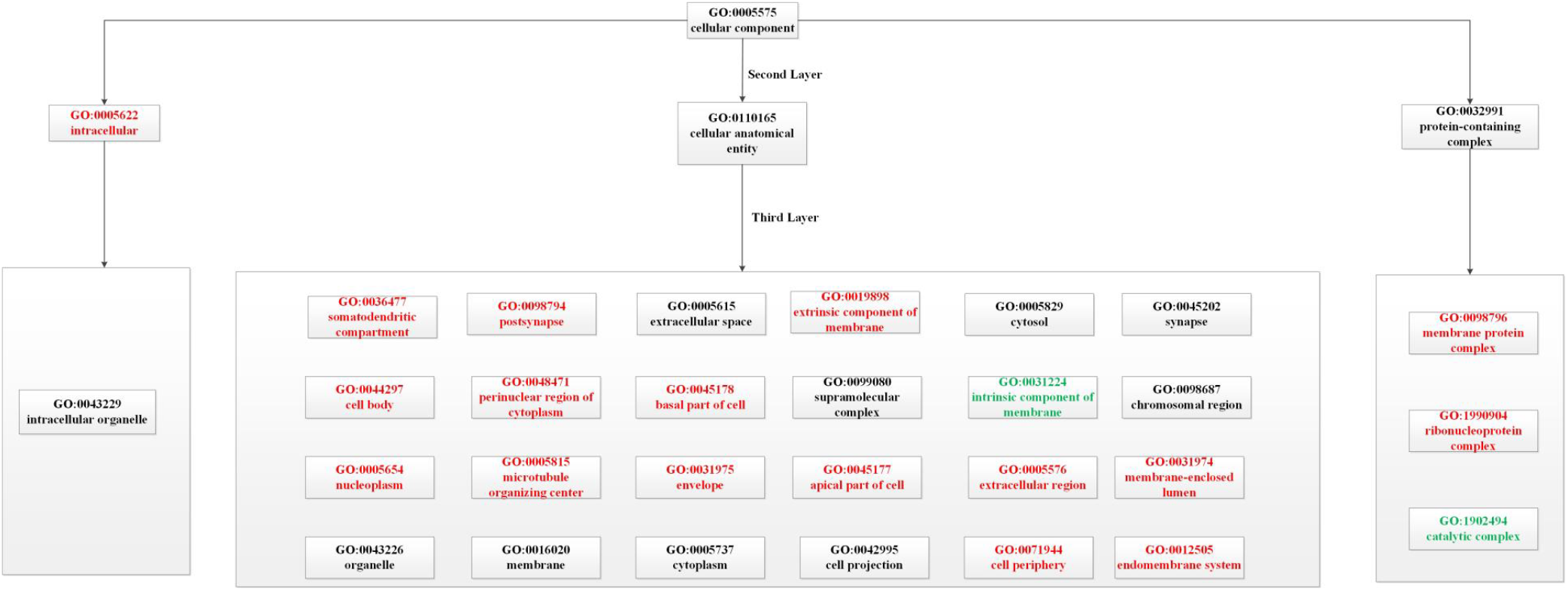
The performance over M-AUPR in CCO. There are 31 GO terms in the second and third layers of the CCO domain that are annotated with more than ten proteins in the test set. By checking the performance improvement of NetGO 3.0 over 2.0 on each term, the GO terms that have more than a 5% increase or decrease are shown in red and green, respectively.

### Performance comparison on difficult proteins

Following the CAFA setting, proteins with a BLAST identity of less than 0.6 to any protein in training data are identified as “difficult proteins” [3]. In the test set, there are 66, 85, and 70 difficult proteins in MFO, BPO, and CCO, respectively. It is evident that methods based on homology find it hard to predict the function of difficult proteins accurately. **Table 5** shows the performance of different methods in dealing with difficult proteins. As mentioned above, BLAST-KNN, a method that annotated target proteins by homology proteins, ranked last in 9 experimental settings. We find that LR-InterPro and LR-ESM are the two best-performing component methods in this scenario. For example, in terms of *S_min_*, there is a slight difference between the two methods in three domains. LR-ESM and LR-InterPro achieve the best performance for all component methods in 6 and 3 out of 9 settings. Once again, NetGO 3.0 is proved to be the best method for predicting the function of difficult proteins.

**Table 5.**
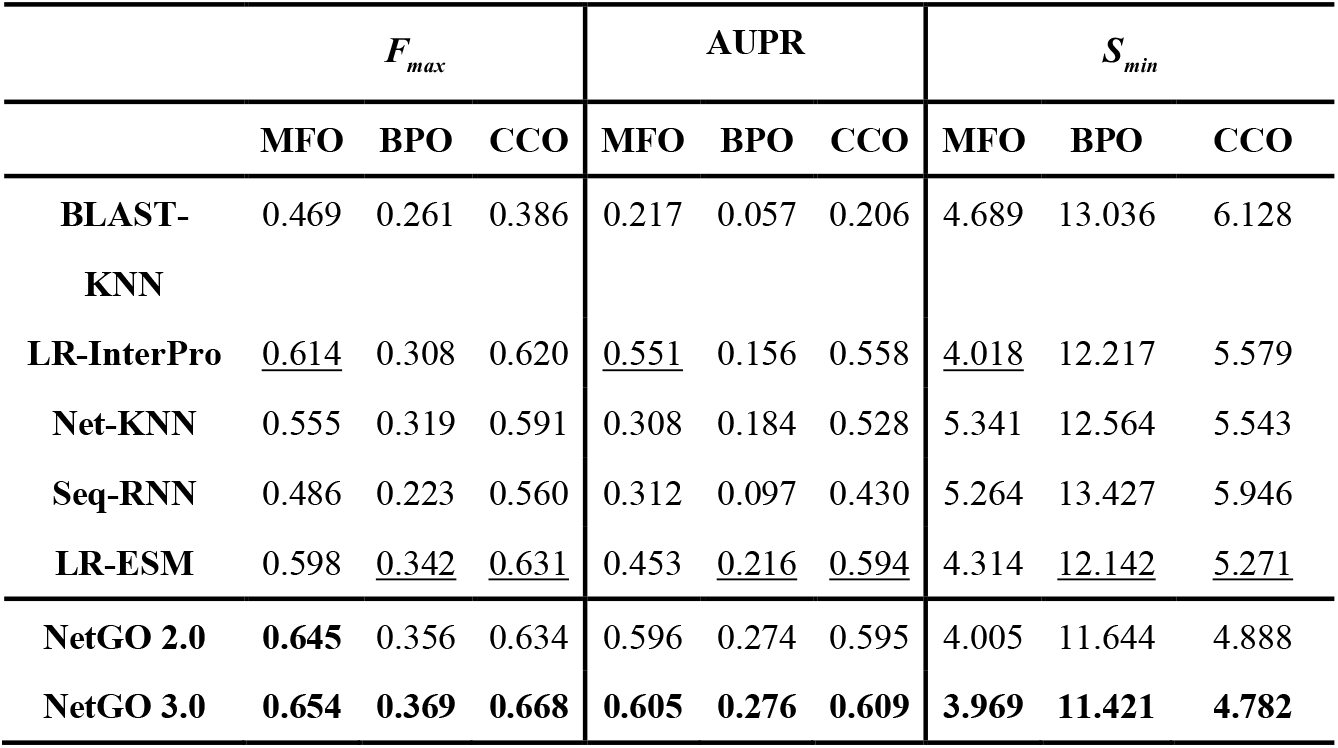
Performance on difficult proteins.

### Performance comparisons on proteins with sequence lengths longer than 1000

We performed a truncation operation for proteins longer than 1000 so that ESM-1b could generate representations for all proteins in the dataset. Focusing on the performance of each method on these long proteins helps us better understand the advantages and limitations of NetGO 3.0. There exist 21, 78, and 26 test proteins in MFO, BPO, and CCO. **Table 6** lists the prediction results of component methods, NetGO 2.0 and NetGO 3.0. As shown in Table 6, LR-ESM is no longer one of the best-performing component methods, which indirectly causes NetGO 3.0 to perform worse than NetGO 2.0 in MFO and BPO. By comparing each method’s performance on the entire test set in Table 1, we noticed that the performance decreased for all methods except Net-KNN. This suggests that function prediction for long proteins is a challenge.

**Table 6.** Performance comparisons on proteins with sequence lengths longer than 1000.

| | $F_{max}$ | | | AUPR | | | $S_{min}$ | | |
| --- | --- | --- | --- | --- | --- | --- | --- | --- | --- |
|  | MFO | BPO | CCO | MFO | BPO | CCO | MFO | BPO | CCO |
| **BLAST-KNN** | 0.514 | 0.272 | 0.549 | 0.271 | 0.119 | 0.465 | 6.349 | 15.176 | 7.072 |
| **LR-InterPro** | <u>0.595</u> | 0.312 | <u>0.638</u> | 0.407 | 0.111 | <u>0.603</u> | <u>5.389</u> | <u>15.454</u> | <u>6.253</u> |
| **Net-KNN** | 0.515 | <u>0.329</u> | 0.609 | <u>0.455</u> | <u>0.205</u> | 0.598 | 6.271 | 14.510 | 6.907 |
| **Seq-RNN** | 0.509 | 0.304 | 0.587 | 0.329 | 0.162 | 0.508 | 6.103 | 14.965 | 6.903 |
| **LR-ESM** | 0.536 | 0.309 | 0.586 | 0.424 | 0.135 | 0.563 | 5.855 | 15.213 | 6.662 |
| **NetGO 2.0** | **0.587** | **0.357** | 0.625 | **0.497** | **0.241** | 0.589 | **5.312** | **13.824** | 6.240 |
| **NetGO 3.0** | 0.577 | 0.348 | **0.631** | 0.485 | 0.215 | **0.606** | 5.452 | 13.947 | **5.938** |

Moreover, we observed the prediction performance of NetGO 2.0 and NetGO 3.0 on several unannotated proteins Q3UZV7, F1QKQ1, and Q2HX28. The sequence lengths of these three proteins are 1028, 1356, and 1409. As shown in **Table 7**, NetGO 2.0 achieved better AUPR on three proteins, which indicates that the truncated sequences in long proteins are important sources of information and are critical for predicting functions. This further confirms that NetGO 3.0 needs to be improved in handling long sequences, which will be important future research work.

**Table 7.**
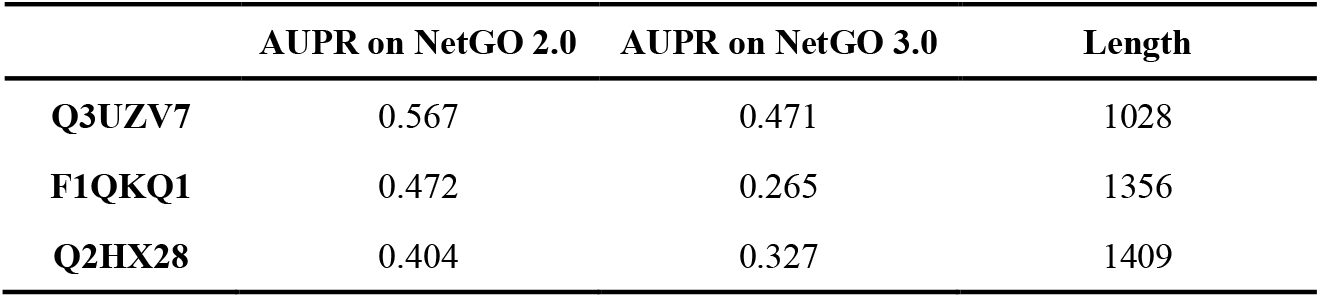
Performance on three long proteins in BPO.

|  | AUPR on NetGO 2.0 | AUPR on NetGO 3.0 | Length |
| --- | --- | --- | --- |
| **Q3UZV7** | 0.567 | 0.471 | 1028 |
| **F1QKQ1** | 0.472 | 0.265 | 1356 |
| **Q2HX28** | 0.404 | 0.327 | 1409 |

### Visualization of the predicted results

We present more options to visualize predicted GO terms to better illustrate prediction results. Compared to NetGO 2.0, the new web server offers a novel perspective to present the results, which can provide more relevant information about predicted GO terms. **Figure 4** shows the new result page of NetGO 3.0, which mainly includes three ways to visualize the prediction performance. Although GO terms in top layers usually achieve a higher score and rank higher, NetGO 3.0 clarifies the depth of predicted GO terms, which allows users to find specific GO terms in bottom layers. Note that the colour in the result page and node size in Figure 4.D are determined by the predicted confidence score, which can help users better understand the predicted results in an original view.

**Figure 4.**
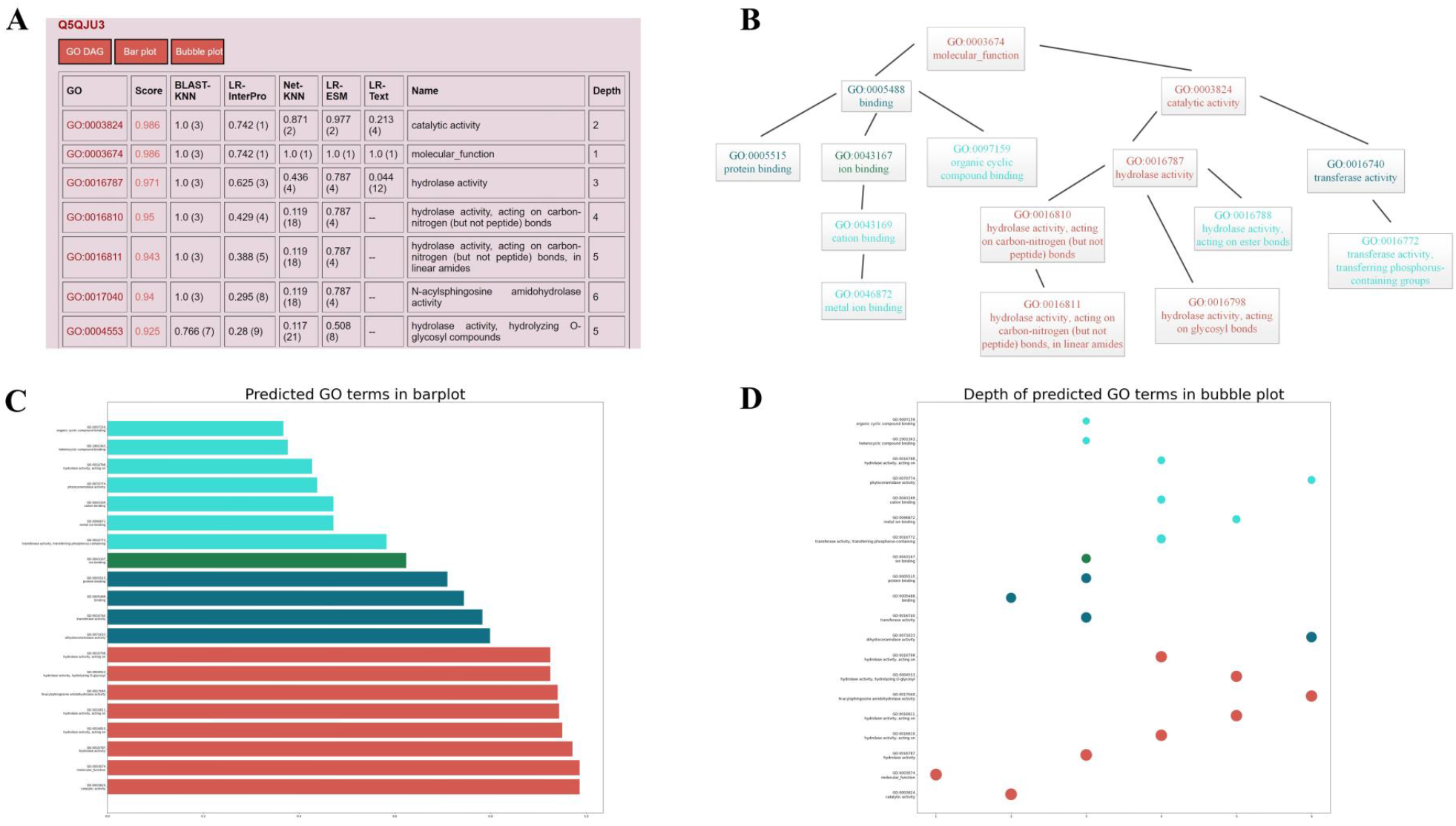
Visualization of prediction results on the web server. A. Prediction result page of NetGO 3.0’s website. “GO DAG”, “Bar plot” and “Bubble plot” are the new interfaces to visualize the predicted GO terms. We also added a new column named “Depth” to show the depth of GO terms in Gene Ontology. B. The predicted GO terms and their directed acyclic graph. C. We plot the predicted GO terms and their confidence scores in a bar diagram. D. The Predicted GO term and its depth in Gene Ontology are plotted in a bubble diagram.

### Case study

Finally, we select a specific protein as input and show the results obtained by NetGO 3.0 and its competing methods. Ubiquitin-like protein 5 (UniProt ID: Q9FGZ9) is a difficult protein with low BLAST similarity to training proteins. **Table 8** shows the 18 GO terms in BPO annotated to protein Q9FGZ9. **Figure 5** also depicts the directed acyclic graphs (DAG) according to the relationship of 18 GO terms in Gene Ontology. In Table 8, it is clear that BLAST-KNN failed to achieve a valid result because homology-based methods were not suitable for difficult protein function prediction. LR-InterPro and LR-ESM extracted features from raw amino acid sequences and obtained better results than BLAST-KNN. In the top 20 predicted GO terms, the number of true positive samples achieved by LR-ESM is significantly larger than other methods, which predicted 14 correct function labels. NetGO and NetGO 2.0 predicted only six correct GO terms, which are not competitive compared to LR-ESM and NetGO 3.0. The reason for this phenomenon may be that the new component method, LR-ESM, is more robust for difficult proteins than other methods and is able to represent them more efficiently. With the support of the protein language model, NetGO 3.0 achieved 15 true GO terms out of 19 predicted ones, which successfully predicted the GO terms that NetGO and NetGO 2.0 failed to predict. Figure 5 illustrates the hierarchy of correct and predicted GO terms, indicating that NetGO 3.0 is able to predict those GO terms with less information in the deeper layers. Overall, this typical example demonstrates that the high predictive performance of NetGO 3.0 is closely related to the protein language models.

**Figure 5.**
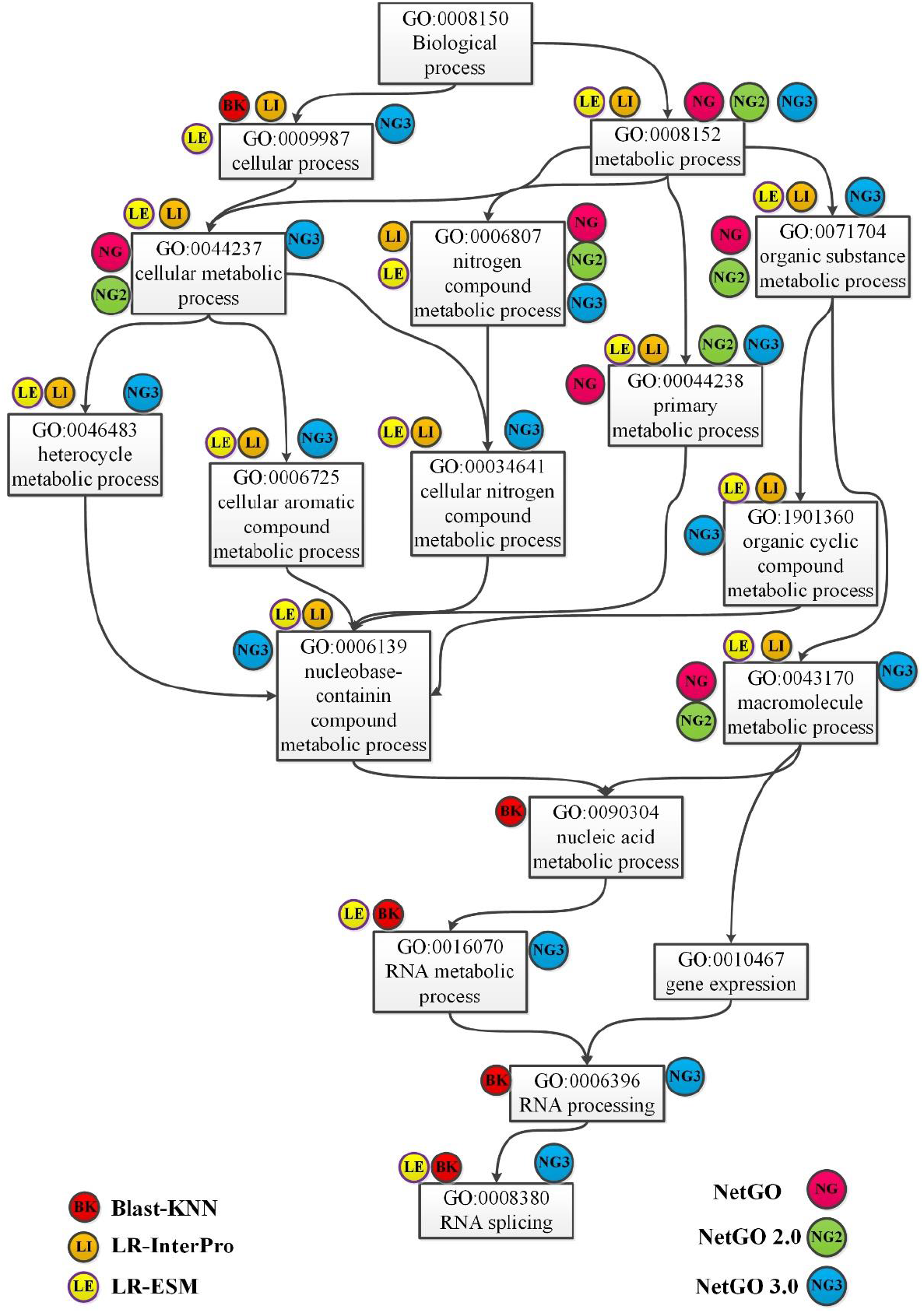
DAG of GO terms associated with Q9FGZ9 in BPO. Each GO term is attached with tags, which illustrates that the GO term is predicted correctly by corresponding methods.

**Table 8.** Predicted GO terms of Q9FGZ9 in BPO by NetGO 3.0 and competing methods. Each method shows the top 20 predicted GO terms (the root term GO:0008150 is deleted). Correctly predicted GO terms are in red, and the last row shows the groundtruth GO terms.

| Method | GO terms |  |  |  |
| --- | --- | --- | --- | --- |
| BLAST-KNN | GO:0009987 | GO:0010628 | GO:0010604 | GO:0010468 |
|  | GO:0060255 | GO:0009893 | GO:0019222 | GO:0048518 |
|  | GO:0050789 | GO:0065007 | GO:0000398 | GO:0000377 |
|  | GO:0000375 | GO:0008380 | GO:0006397 | GO:0016071 |
|  | GO:0006396 | GO:0016070 | GO:0090304 |  |
| LR-InterPro | GO:0009987 | GO:0008152 | GO:0050896 | GO:0044237 |
|  | GO:0071704 | GO:0044238 | GO:0006807 | GO:0043170 |
|  | GO:0065007 | GO:0050789 | GO:0006950 | GO:0044260 |
|  | GO:1901564 | GO:0034641 | GO:0044267 | GO:0019538 |
|  | GO:1901360 | GO:0006725 | GO:0046483 |  |
| LR-ESM | GO:0009987 | GO:0043170 | GO:0071704 | GO:0008152 |
|  | GO:0044238 | GO:0006807 | GO:0044260 | GO:0044237 |
|  | GO:0050896 | GO:0034641 | GO:0046483 | GO:1901360 |
|  | GO:0006139 | GO:0006725 | GO:0043412 | GO:0008380 |
|  | GO:0016070 | GO:0000375 | GO:000377 |  |
| NetGO | GO:0009987 | GO:0008152 | GO:0044237 | GO:0050896 |
|  | GO:0043170 | GO:0071704 | GO:0006807 | GO:0044238 |
|  | GO:0048856 | GO:0032502 | GO:0032501 | GO:0006950 |
|  | GO:0050789 | GO:0065007 | GO:0007275 | GO:0000003 |
|  | GO:0022414 | GO:0051704 | GO:0042221 |  |
| NetGO 2.0 | GO:0009987 | GO:0044237 | GO:0008152 | GO:0050896 |
|  | GO:0071704 | GO:0044238 | GO:0050789 | GO:0065007 |
|  | GO:0043170 | GO:0006807 | GO:0032502 | GO:0048856 |
|  | GO:0006950 | GO:0050794 | GO:0007275 | GO:0032501 |
|  | GO:0003006 | GO:0022414 | GO:0000003 |  |
| NetGO 3.0 | GO:0009987 | GO:0008152 | GO:0044237 | GO:0044238 |
|  | GO:0043170 | GO:0071704 | GO:0006807 | GO:0050896 |
|  | GO:0050789 | GO:0065007 | GO:1901360 | GO:0046483 |
|  | GO:0006139 | GO:0006725 | GO:0034641 | GO:0006950 |
|  | GO:0008380 | GO:0006396 | GO:0016070 |  |
| Groudtruth | GO:0044237 | GO:0044238 | GO:0009987 | GO:0006139 |
|  | GO:0046483 | GO:0008150 | GO:0006807 | GO:0071704 |
|  | GO:0006396 | GO:0016070 | GO:0010467 | GO:0008380 |
|  | GO:1901360 | GO:0008152 | GO:0090304 | GO:0043170 |
|  | GO:0034641 | GO:0006725 |  |  |

## Conclusion

We have developed NetGO 3.0 to improve the performance of large-scale AFP by incorporating a new component LR-ESM, which utilizes a protein language model to generate powerful representations of proteins. Interesting future work would be integrating protein structural information into NetGO to enhance the performance of AFP [18–20].

## Availability

The web server of NetGO 3.0 is freely accessible at https://dmiip.sjtu.edu.cn/ng3.0.

## CRediT author statement

**Shaojun Wang:** Data Curation, Software, Methodology, Writing - original draft. **Ronghui You:** Conceptualization, Methodology, Writing - review & editing. **Yunjia Liu:** Methodology, Visualization, Writing - review & editing. **Yi Xiong:** Resource, Writing - review & editing. **Shanfeng Zhu:** Conceptualization, Resource, Writing – review & editing. All authors read and approved the final manuscript.

## Acknowledgments

S.Z. has been supported by National Natural Science Foundation of China (No. 61872094), Shanghai Municipal Science and Technology Major Project (No.2018SHZDZX01), ZJ Lab, and Shanghai Center for Brain Science and Brain-Inspired Technology. S.W and R.Y. have been supported by the 111 Project (No. B18015), Shanghai Municipal Science and Technology Major Project (No. 2017SHZDZX01) and Information Technology Facility, CAS-MPG Partner Institute for Computational Biology, Shanghai Institute for Biological Sciences, Chinese Academy of Sciences. We are thankful to Prof. Huang and Ms. Sarah Replogle for English proofreading.

## Supplementary material

**Section S1 Definition of performance evaluation metrics**

**Section S2 Details of the new dataset**

**Section S3 Performance of different competing models**

**Table S1 Summary of the benchmark dataset**

**Table S2 Summary of the new dataset**

**Table S3 Performance of different competing models**

## References

[1] Ashburner M, Ball C A, Blake J A, Botstein D, Butler H and Cherry J M, et al. Gene Ontology: tool for the unification of biology. Nat Genet 2000;25:25–9.

[2] Consortium T U. UniProt: the universal protein knowledgebase in 2021. Nucleic Acids Res 2021;49:D480–9.

[3] Zhou N, Jiang Y, Bergquist T R, Lee A J, Kacsoh B Z and Crocker A W, et al. The CAFA challenge reports improved protein function prediction and new functional annotations for hundreds of genes through experimental screens. GENOME BIOL 2019;20:1–23.

[4] Piovesan D and Tosatto S C. INGA 2.0: improving protein function prediction for the dark proteome. Nucleic Acids Res 2019;47:W373–8.

[5] Kulmanov M and Others. DeepGOWeb: fast and accurate protein function prediction on the (Semantic) Web. Nucleic Acids Res 2021;49:W140–6.

[6] Zhang C, Zheng W, Freddolino P L and Zhang Y. MetaGO: Predicting Gene Ontology of Non-homologous Proteins Through Low-Resolution Protein Structure Prediction and Protein–Protein Network Mapping. J MOL BIOL 2018;430:2256–65.

[7] Smaili F Z, Tian S, Roy A, Alazmi M, Arold S T and Mukherjee S, et al. QAUST: Protein Function Prediction Using Structure Similarity, Protein Interaction, and Functional Motifs. Genomics, Proteomics & Bioinformatics 2021;

[8] Li H. A Short Introduction to Learning to Rank. IEICE Transactions 2011;94-D(10):1854–62.

[9] You R, Zhang Z, Xiong Y, Sun F, Mamitsuka H and Zhu S. GOLabeler: improving sequence-based large-scale protein function prediction by learning to rank. Bioinformatics 2018;34:2465–73.

[10] You R, Yao S, Xiong Y, Huang X, Sun F and Mamitsuka H, et al. NetGO: improving large-scale protein function prediction with massive network information. Nucleic Acids Res 2019;47:W379–87.

[11] Yao S, You R, Wang S, Xiong Y, Huang X and Zhu S. NetGO 2.0: improving large-scale protein function prediction with massive sequence, text, domain, family and network information. Nucleic Acids Res 2021;49:W469–75.

[12] Devlin J, Chang M, Lee K and Toutanova K. BERT: Pre-training of Deep Bidirectional Transformers for Language Understanding. NAACL-HLT 2019;4171–86.

[13] Rives A, Meier J, Sercu T, Goyal S, Lin Z and Liu J, et al. Biological structure and function emerge from scaling unsupervised learning to 250 million protein sequences. Proceedings of the National Academy of Sciences 2021;118:e2016239118.

[14] Elnaggar A, Heinzinger M, Dallago C, Rehawi G, Wang Y and Jones L, et al. ProtTrans: Towards Cracking the Language of Lifes Code Through Self-Supervised Deep Learning and High Performance Computing. TPAMI 2021;44(10).

[15] Alley E C, Khimulya G, Biswas S, AlQuraishi M and Church G M. Unified rational protein engineering with sequence-based deep representation learning. Nat Methods 2019;16:1315–22.

[16] Strodthoff N, Wagner P, Wenzel M and Samek W. UDSMProt: universal deep sequence models for protein classification. Bioinformatics 2020;36:2401–9.

[17] Villegas-Morcillo A, Makrodimitris S, van Ham R C, Gomez A M, Sanchez V and Reinders M J. Unsupervised protein embeddings outperform hand-crafted sequence and structure features at predicting molecular function. Bioinformatics 2021;37:162–70.

[18] Lai B and Xu J. Accurate protein function prediction via graph attention networks with predicted structure information. Brief in Bioinform 2022;23(1):bbab502.

[19] Zhang C, Freddolino P L and Zhang Y. COFACTOR: improved protein function prediction by combining structure, sequence and protein--protein interaction information. Nucleic Acids Res 2017;45:W291–9.

[20] Gligorijević V, Renfrew P D, Kosciolek T, Leman J K, Berenberg D, and Vatanen T, et al. Structure-based protein function prediction using graph convolutional networks. Nat Commun 2021;12:1–14.

[21] Rao R, Bhattacharya N, Thomas N, Duan Y, Chen X, and Canny J, et al. Evaluating protein transfer learning with TAPE. NeurIPS 2019;32:9686–98.

[22] Bepler T and Berger B. Learning the protein language: Evolution, structure, and function. Cell Syst 2021;12:654–69.

[23] Huntley R P, Sawford T, Mutowo-Meullenet P, Shypitsyna A, Bonilla C and Martin MJ, et al. The GOA database: gene Ontology annotation updates for 2015. Nucleic Acids Res 2015;43:D1057–63.

