## Supplemental text for "NetGO 3.0: Protein Language Model Improves Large-scale Functional Annotations"

**Supplementary material**

**SectionS1 Definition of performance evaluation metrics**

AUPR,
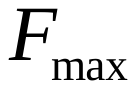
 and
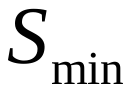
 are three main metrics to evaluate the performance. AUPR is a classical evaluation metric in machine learning especially for imbalanced classification, which punishes false positive prediction.
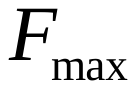
 is an official metric of CAFA with the following definition.

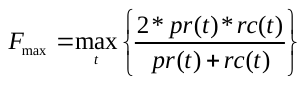
(1)

where pr(t) and rc(t) are precision and recall, respectively, obtained at a cut-off value t, which is defined as follows:

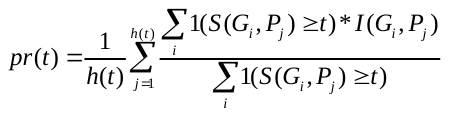
(2)

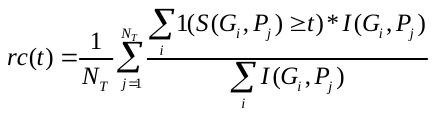
(3)

where h(t) is the number of proteins with the score no smaller than t for at least one GO term.

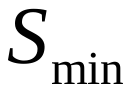
 means the minimum semantic distance with definition as follows:

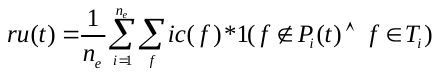
(4)

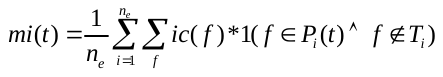
(5)

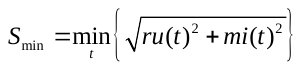
(6)

where ru(t) and mi(t) denote remaining uncertainty and misinformation at certain threshold t. In the above formulas, ic calculates the information content of GO term, given as follows:

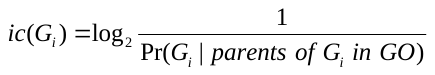
(7)

where Pr(G_i_ | parents of G_i_ in GO) is the conditional probability of G_i_ given its parents of the GO structure.

**Section S2 Details of new dataset**

NetGO 3.0 collects data by following the procedures of CAFA, which mainly focus two types of proteins: no-knowledge and limited-knowledge proteins. In NetGO 3.0, no-knowledge proteins do not have any experimental annotations before 2020, while limited-knowledge proteins have at least one annotation in the other domains before 2020. Table S2 lists the numbers of proteins in the dataset.

1. Training data: all experimental annotation data before January 2020.
2. Validation data: all experimental *no-knowledge* and *limited-knowledge* proteins annotated from January 2020 to December 2020.
3. Testing data: all experimental *no-knowledge* proteins between January 2021 and December 2021.

**Section S3 Performance of different competing models**

As listed in Table S3, simply adding LR-ESM into NetGO 2.0 only makes a slight improvement compared with NetGO 3.0. Two classifiers are evenly matched, each with its strengths. NetGO 2.0 with LR-ESM performs better in BPO and CCO while NetGO 3.0 has a slight advantage in MFO. However, the difference is subtle. Considering computing resources, complexity and other factors, we replace Seq-RNN with LR-ESM as a component method to complete NetGO 3.0.

**Table S1 Summary of benchmark dataset**

**Table S1 Summary of new dataset**

**Table S3 Performance of different competing models**
