## Supplemental Table 1 for "NetGO 3.0: Protein Language Model Improves Large-scale Functional Annotations"

**Table S1 Summary of benchmark dataset**

|  | **Train** | | | **LTR** | | | **Test** | | |
| --- | --- | --- | --- | --- | --- | --- | --- | --- | --- |
|  | **MFO** | **BPO** | **CCO** | **MFO** | **BPO** | **CCO** | **MFO** | **BPO** | **CCO** |
| **HUMAN** | 9315 | 12492 | 19014 | 232 | 183 | 0.464 | 194 | 45 | 50 |
| **MOUSE** | 6316 | 10355 | 8879 | 209 | 213 | 0.405 | 37 | 74 | 53 |
| **DROME** | 5295 | 10084 | 7223 | 219 | 192 | 0.478 | 30 | 49 | 20 |
| **ARATH** | 5191 | 8784 | 9664 | 451 | 248 | 0.587 | 91 | 47 | 36 |
| **DANRE** | 2606 | 11082 | 2057 | 148 | 291 | 0.528 | 32 | 176 | 17 |
| **RAT** | 4360 | 5538 | 5096 | 35 | 57 | 0.477 | 13 | 21 | 29 |
| **ALL** | 52923 | 88060 | 78842 | 1841 | 1546 | 1747 | 444 | 491 | 264 |
