## Supplemental Table 2 for "NetGO 3.0: Protein Language Model Improves Large-scale Functional Annotations"

**Table S1 Summary of new dataset**

|  | **Train** | | | **LTR** | | | **Test** | | |
| --- | --- | --- | --- | --- | --- | --- | --- | --- | --- |
|  | **MFO** | **BPO** | **CCO** | **MFO** | **BPO** | **CCO** | **MFO** | **BPO** | **CCO** |
| **HUMAN** | 9380 | 11181 | 19218 | 641 | 131 | 941 | 49 | 66 | 61 |
| **MOUSE** | 6412 | 10493 | 9005 | 273 | 224 | 189 | 64 | 119 | 95 |
| **DROME** | 5404 | 10045 | 7340 | 170 | 229 | 166 | 107 | 169 | 152 |
| **ARATH** | 5071 | 8832 | 8469 | 839 | 230 | 179 | 73 | 129 | 127 |
| **DANRE** | 2743 | 11260 | 2089 | 228 | 374 | 71 | 10 | 105 | 13 |
| **RAT** | 4236 | 5414 | 5041 | 41 | 56 | 73 | 17 | 47 | 31 |
| **ALL** | 53680 | 87093 | 78458 | 2491 | 1654 | 1866 | 423 | 816 | 625 |
