## Supplemental Table 3 for "NetGO 3.0: Protein Language Model Improves Large-scale Functional Annotations"

**Table S3 Performance of different competing models**

|  | 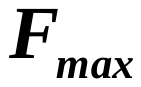 | | | **AUPR** | | | 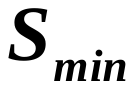 | | |
| --- | --- | --- | --- | --- | --- | --- | --- | --- | --- |
|  | **MFO** | **BPO** | **CCO** | **MFO** | **BPO** | **CCO** | **MFO** | **BPO** | **CCO** |
| **DeepGOWeb** | 0.620 | 0.605 | 0.620 | 0.521 | 0.115 | 0.493 | 4.496 | 14.772 | 5.550 |
| **GOLabeler** | 0.667 | 0.326 | 0.631 | 0.647 | 0.193 | 0.557 | 3.970 | 13.558 | 5.295 |
| **NetGO 2.0** | 0.666 | 0.366 | 0.663 | 0.655 | 0.269 | 0.593 | 4.013 | 12.984 | 4.756 |
| **NetGO 2.0 +LR-ESM** | **0.680** | 0.377 | **0.670** | **0.673** | **0.270** | 0.619 | 3.841 | **12.797** | **4.691** |
| **NetGO 3.0** | 0.679 | **0.378** | **0.670** | 0.672 | 0.268 | **0.620** | **3.840** | 12.800 | 4.735 |
